# Differentiation of Bacterial, Fungal, and Algal Communities on Coastal Concrete Versus Drainage Pipes and the Coexistence of Corrosion and Healing Potentials

**DOI:** 10.64898/2026.06.03.729873

**Authors:** Shengxun Yao, Maomi Zhao, Jing Xiang, Xiufen Liao, Qiumei Jiang, Congtao Sun, Yan Wang

## Abstract

Coastal concrete structures and drainage pipes are prone to microbially influenced deterioration. However, differences in microbial communities and their corrosion/healing potentials between these habitats remain unclear. Here, we compared bacterial(16S), fungal (ITS) and algal(18S) communities on coastal concrete(C) and drainage pipe(P) surfaces. Fungal and algal α-diversity were significantly higher in P than in C, while bacterial diversity did not differ. β-Diversity strongly separated bacterial and algal communities between habitats, but not fungi. A shared core “seed bank” of 575 bacterial, 520 fungal and 40 algal ASVs was identified. Student’s *t-*test revealed that P enriched oligotrophic degraders (*Sphingomonas*) and acid-producing fungi (*Arxiella*, *Bisifusarium*), whereas C selected for halotolerant EPS-producing bacteria (*Tunicatimonas*, *Muricauda*) and the extremotolerant alga *Coelastrella*. db-RDA linked these differences to salinity, NH_4_^+^-N, NO_3_^-^-N, and COD. Functional prediction indicated a shift from metabolism pathways in C to signaling in P. Co-occurrence networks revealed cross-kingdom competition and within-kingdom cooperation, especially among algae. Importantly, both habitats harbored microorganisms with documented corrosion and healing potentials, but under natural conditions, net deterioration dominated, microbial healing is hardly to counteract the negative effects. This study provides a functional taxonomic framework for understanding and managing concrete microbiomes in coastal and sewer infrastructure.

**Importance:** Concrete is the most widely used construction material, and its deterioration in coastal and sewer environments poses significant economic and safety challenges. Microorganisms play a dual role in concrete durability — they can both corrode and heal concrete, but the net outcome under natural conditions is poorly understood. Most studies have focused on bacteria alone, overlooking the contributions of fungi and algae, and few reports on the of coastal concrete which also exist the microbially influenced concrete corrosion (MICC) similar to sewer. Here, we simultaneously analyzed all three microbial kingdoms on coastal concrete and drainage pipes. We found that while concrete surfaces harbor a diverse “seed bank” of microorganisms potentially involved in both corrosion and healing, under natural conditions, net deterioration dominated, indicating microbial healing is hard to counteract the negative effects from the environment and microbial. Our work provides a functional framework to guide the development of microbiome-based strategies for enhancing concrete durability, such as activating rare healing taxa in drainage pipes or selecting for endogenous healers in marine environments.

## 1 Introduction

Concrete is the most widely used construction material globally, and its long-term durability is critical for infrastructure sustainability (1). In coastal and sewer environments, concrete structures are exposed to aggressive chemical and biological attacks that accelerate deterioration. Microbially influenced concrete corrosion (MICC) is a particularly severe problem, leading to substantial economic losses and safety risks (2). In sewer systems, hydrogen sulfide generated by sulfate-reducing bacteria is oxidized to sulfuric acid by sulfur-oxidizing bacteria (SOB), which dissolves cement hydration products and forms expansive gypsum, resulting in rapid material loss (3), (4). In marine environments, high salinity, tidal fluctuations, and biofilm formation further exacerbate concrete degradation(1). Understanding the microbial processes driving such deterioration is therefore essential for developing effective protection strategies and extending the service life of concrete infrastructure.

The composition and assembly of microbial communities on concrete surfaces have been increasingly characterized using modern molecular techniques. Early culture-dependent studies identified key bacterial groups on corroded concrete, including *Acidithiobacillus*, *Thiobacillus*, and *Mycobacterium* (5). With the advent of high-throughput sequencing, the complexity of concrete-associated microbiomes has become more apparent. In sewer systems, microbial communities exhibit pronounced spatial heterogeneity: the upper parts of pipes are dominated by sulfur-oxidizing bacteria such as *Acidithiobacillus* and *Thiothrix*, while the bottom parts harbor sulfate-reducing bacteria like *Desulfovibrio* and *Desulfobulbus* as well as fermentative bacteria (6). Environmental factors including pH, moisture, hydrogen sulfide concentration, and organic carbon strongly shape these communities (7). In marine environments, concrete surfaces develop biofilms rich in Proteobacteria, Cyanobacteria, and Bacteroidetes, with successional patterns influenced by surface material and exposure time (1, 8). Coastal conditions impose unique selective pressures, favoring halotolerant and desiccation-resistant taxa such as *Rubrobacter* and *Tunicatimonas* (1, 9). However, most studies have focused solely on bacteria, neglecting the important roles of fungi and algae. Filamentous fungi, including *Aspergillus*, *Penicillium*, *Cladosporium*, and *Fusarium*, are frequently isolated from damp concrete and mortar, where they produce organic acids and physically penetrate the matrix (10, 11). Algae, particularly green macroalgae like *Ulva*, *Chaetomorpha*, and *Rhizoclonium*, and diatoms like *Navicula* and *Achnanthes*, colonize moist surfaces, extract calcium for metabolism, and form biofilms that alter local chemistry (12, 13). Despite these insights, systematic research on bacteria, fungi and algae across different concrete substrates, especially coastal concrete and drainage pipes, remains lacking.

The microbial mechanisms driving concrete corrosion have been extensively investigated, primarily focusing on sulfur-cycling bacteria. *Acidithiobacillus thiooxidans* is recognized as a key acidophilic SOB that produces large amounts of sulfuric acid, causing severe concrete deterioration (2, 3). Other SOB, such as *Thiothrix* and *Halothiobacillus*, contribute to the early and intermediate stages of corrosion (3). Sulfate-reducing bacteria like *Desulfovibrio* and *Desulfobulbus* could supply H_2_S to SOB, completing the sulfur cycle that drives corrosion (6). Fungi are increasingly acknowledged as active concrete deteriorators. *Fusarium oxysporum*, *Aspergillus niger*, and *Cladosporium sphaerospermum* secrete organic acids like oxalic, citric, and gluconic that chelate calcium and dissolve cement hydrates; their hyphae can penetrate up to 620 µm into the concrete matrix (10, 14). Algae also contribute to biodeterioration include macroalgae such as *Ulva fasciata* and *Chaetomorpha antennina* cause substantial surface damage through calcium uptake and biofilm-mediated pH changes(12, 13, 15). However, most previous studies have examined individual microbial groups in isolation, limiting our understanding of multi-kingdom interactions and their combined effects on concrete durability.

Beyond deterioration, concrete-associated microorganisms possess self-healing potential hrough microbially induced calcium carbonate precipitation (MICP). Extensive research has demonstrated that ureolytic bacteria, particularly *Bacillus pasteurii*, *B. subtilis*, *B. megaterium*, and *Sporosarcina pasteurii*, can precipitate calcite and effectively seal cracks (16–18). Other bacteria, such as *Exiguobacterium mexicanum* and *Pseudomonas spp*., also exhibit MICP activity under saline or nutrient-rich conditions (9, 19). More recently, fungi have been explored as self-healing agents. *Aspergillus nidulans*, *Penicillium chrysogenum*, *Trichoderma reesei*, and *Fusarium oxysporum* can induce calcium carbonate biomineralization via urease or non-ureolytic pathways (20–22). Even yeasts like *Candida orthopsilosis* and *Rhodotorula mucilaginosa* have been shown to form vaterite (23). Notably, many genera like *Aspergillus*, *Penicillium*, *Fusarium*, *Bacillus*, and *Pseudomonas* contain both corrosive and healing species, making the net outcome highly context-dependent (10, 22). This functional duality underscores the need to simultaneously evaluate both deteriorative and reparative potentials within natural concrete microbiomes.

Despite the growing body of research, key gaps remain. In particular, how microbial communities differentiate between coastal concrete and drainage pipe surfaces, and how corrosion-related and healing-related potentials coexist within these communities, have not been systematically addressed.

To fill these gaps, we compared bacterial, fungal, and algal communities on coastal concrete surfaces (C) and drainage concrete pipes (P) using 16S, ITS, and 18S amplicon sequencing. Our objectives were to characterize the diversity and composition of the three microbial kingdoms on each substrate; identify differential abundant taxa and link them to environmental factors like salinity, NH_4_^+^-N, NO_3_^-^-N, and COD; predict functional pathways of bacterial communities and construct cross-domain co-occurrence networks; and catalog potential corrosion-related and healing-related microorganisms based on literature, and evaluate their distribution strategies. We hypothesized that environmental filtering drives distinct microbial assemblages between C and P, and that although both corrosion and healing potentials coexist, the net outcome under natural conditions is dominated by deterioration. This study provides a functional taxonomic framework for understanding the dual roles of concrete-associated microbiomes and to supports the development of microbiome-based durability management strategies for marine and sewer infrastructure.

## 2 Method

### 2.1 Sample collection and seawater/water physicochemical indicator measurement

A total of 44 samples includes 16 biofilm samples from coastal concrete (C) and 6 biofilm samples from drainage concrete pipes (P), 16 seawater samples from the sea surface around the coastal concrete, and 6 water samples from the drainage pipes were collected along the coasts of Beihai and Fangcheng City in August 2025. The biofilm samples were immediately placed on ice in the field, transported to the laboratory, and ultimately stored at -80°C until further analysis. Water samples were used for physicochemical analyses. Temperature, dissolved oxygen (DO), and pH were measured in situ using a JPBJ-608 portable meter (INESA, China) for DO and a PHBJ-261L pH meter (INESA, China) for pH. Salinity was determined with a DLX-ARH100 salinometer (DEILIXI, China). Concentrations of NH_4_ ^+^-N and NO_3_ ^-^-N (San++, Skalar, Dutch) were measured in the laboratory using a San⁺⁺ continuous flow analyzer (Skalar, Netherlands). Chemical oxygen demand (COD) was determined by the dichromate digestion method (GB 11914-89) with a detection limit of 2 mg/L.

### 2.2 DNA Extraction, PCR Amplification and Library Construction

Total microbial genomic DNA was extracted using the E.Z.N.A.® soil DNA Kit (Omega Bio-tek, Norcross, GA, U.S.) according to the manufacturer’s instructions. DNA purity and concentration were assessed using a NanoDrop ND-2000 spectrophotometer (Thermo Fisher, USA), and the DNA was stored at −80 ℃ until further use.

The V3-V4 hypervariable region of the 16S rRNA gene was amplified using the primer pair 338F (5’-ACTCCTACGGGAGGCAGCAG -3’) and 806R (5’-GGACTACHVGGGTWTCTAAT-3’), with random 6-base oligo barcode labelling (^24^), the ITS gene using the primer pairs ITS3F (5’-GCATCGATGAA GAACGCAGC-3’) and ITS4R (5’-TCCTCCGCTTATTGATATGC-3’) for fungi (25), and the V4 region of the 18S gene using the primer pairs 3NDF (5’-GGCAAGTCTGGTGCCAG-3’) and V4-euk-R2R (5’-ACGGTATCTRATCRTCTTCG-3’) (26) in T100 Thermal Cycler PCR thermocycler (Bio-Rad, USA). The PCR mixture (20 μL) contained 10 μL 2 × Pro T buffer, 2 μL 2.5 mM dNTPs, 0.8 μL each primer (5 μM), 0.4 μL Fast Pfu polymerase, 10 ng of template DNA, and was adjust to 20 μL with ddH_2_O. The amplification conditions were initial denaturation at 95 ℃ for 3 min; followed by 25 cycles of denaturation at 95℃ for 30 s, annealing at 55 ℃ for 30 s, and extension at 72 ℃ for 45 s; and a final extension at 72 ℃ for 10 min. Negative controls (no template) were included in each PCR run and showed no amplification. After purification of the PCR products using Agencourt AMPure XP beads (Beckman Coulter, USA), sequencing libraries were constructed using the NEXTFLEX Rapid DNA-Seq Kit (Bioo Scientific, USA).

### 2.3 Sequencing and Data Processing

Purified amplicons were pooled in equimolar amounts and paired-end sequenced on an Illumina Nextseq 2000 platform (Illumina, San Diego, USA) according to the standard protocols by Majorbio Bio-Pharm Technology Co. Ltd. (Shanghai, China).

Raw sequencing data were processed with fastp (0.23.4) (27) to remove low-quality reads (Q < 20), and the remaining reads were merged using FLASH (v1.2.11) (28). Amplicon sequence variant (ASV) clustering and denoising were performed using the DADA2 plugin within the QIIME2 (2024.5)(29). Bacterial 16S rRNA gene sequences were taxonomically annotated against the SIlVA 138.2 database at a confidence threshold of 0.7. For fungi, the ITS sequences were classified using the UNITE database (v9.0, https://unite.ut.ee/). For algae (18S rRNA), sequences were classified using the PR2 database (v4.14)(30). To account for disparities in sequencing depth, all samples were rarefied to the minimum sequence count. The raw sequencing data generated in this study have been submitted to the NCBI database under BioProject accession number PRJNA1354633.

### 2.4 Bioinformatics and Statistical Analysis

Alpha diversity indices, including the Chao1 richness index and the Shannon diversity index, were calculated using Mothur v1.30.1 based on the ASVs information 8. Principal coordinate analysis (PCoA) was conducted on a Bray – Curtis dissimilarity matrix, with intergroup differences assessed via PERMANOVA (adonis2 function, 999 permutations) implemented in the vegan package (v2.5-3) in R version 4.2.2. Venn diagrams were generated using the VennDiagram v1.7.3 package in R. For comparisons among different groups, the Wilcoxon rank-sum test was used, followed by false discovery rate (FDR) correction. The significance level was set at *P* < 0.05. The distance-based redundancy analysis (db-RDA) was performed to investigate effect of seawater/water physicochemical parameters on microbial community structure using the same vegan package. The co-occurrence networks were constructed using the top 45 most abundant ASVs of bacteria, fungi, and algae in C and P, respectively, and the top 15 most abundant ASVs of the three in C and P based on a Spearman correlation threshold (|*r*| ≥ 0.5, *p* < 0.05). Key topological metrics include average path length, network diameter, average degree, clustering coefficient, and graph density were calculated using the Gephi platform (31). Functional gene prediction of bacterial communities was performed using PICRUSt2 v2.5.0(32) based on the 16S rRNA gene sequences, and functional annotations were referenced against the FAPROTAX database(33).

### 2.6 Microbial abundance and distribution classification methods

To analyze the abundance and distribution of microbial taxa potentially involved in concrete deterioration and concrete healing, we defined that “abundant” taxa were those with a relative abundance of at least 1% in one or more samples; any genus consistently falling below this threshold was classified as “permanently rare”. For both abundance categories, we defined three genus categories based on occurrence: “broad” (present in ≥75% of samples), “intermediate” (present in >10% but <75% of samples), and “narrow” (present in ≤10% of samples)](34).

## 3 Results

### 3.1 Microbial diversity on C and P

A total of 1,587,342 sequencing reads of bacteria, 1,772,735 of fungi and 998810 of algae were obtained, and after filtering the raw data of 22 samples for quality and chimera, and 10,932 ASVs of bacteria, 4,945 ASVs of fungi, and 390 ASVs of algae were identified. All the ASVs were further classified into 43 bacterial phyla, 13 fungal phyla, and 10 algae phyla based on the reads.

To evaluate and compare the microbial diversity between coastal concrete and concrete pipes, α-diversity indices were calculated based on the ASVs relative abundance of each sample, including the Shannon index and the Chao1 index. The α-diversity showed that the bacterial Chao1 index of C (802.22±169.23) was lower than that of P (878.96±203.80), as well as the Shannon index of C (5.62±0.49) compared to P (5.76±0.54), but neither difference was significant (Figure 1A-B). For the fungi, the Chao1 index of C (164.59±74.39) was significantly lower than that of P (795.75±260.29), as well as the Shannon index of C (3.63±0.91) compared to P (4.81±0.56) (Figure 1B-C). As for the algae, the Chao1 index of C (27.67 ± 13.28) was significantly lower than that of P (36.83 ± 4.12), while the Shannon index of C (2.09 ± 0.64) was lower than that of P (2.42 ± 0.52), but this difference was not significant (Figure 1D-E). Collectively, these results indicate that drainage pipes(P) provide a more favorable ecological niche (with higher diversity) for fungi and algae, whereas bacterial communities exhibit stronger cross-matrix adaptability, suggesting that substrate types exerts selective effects on microbial richness and diversity, with the magnitude varying among microbial groups.

**Fig. 1.**
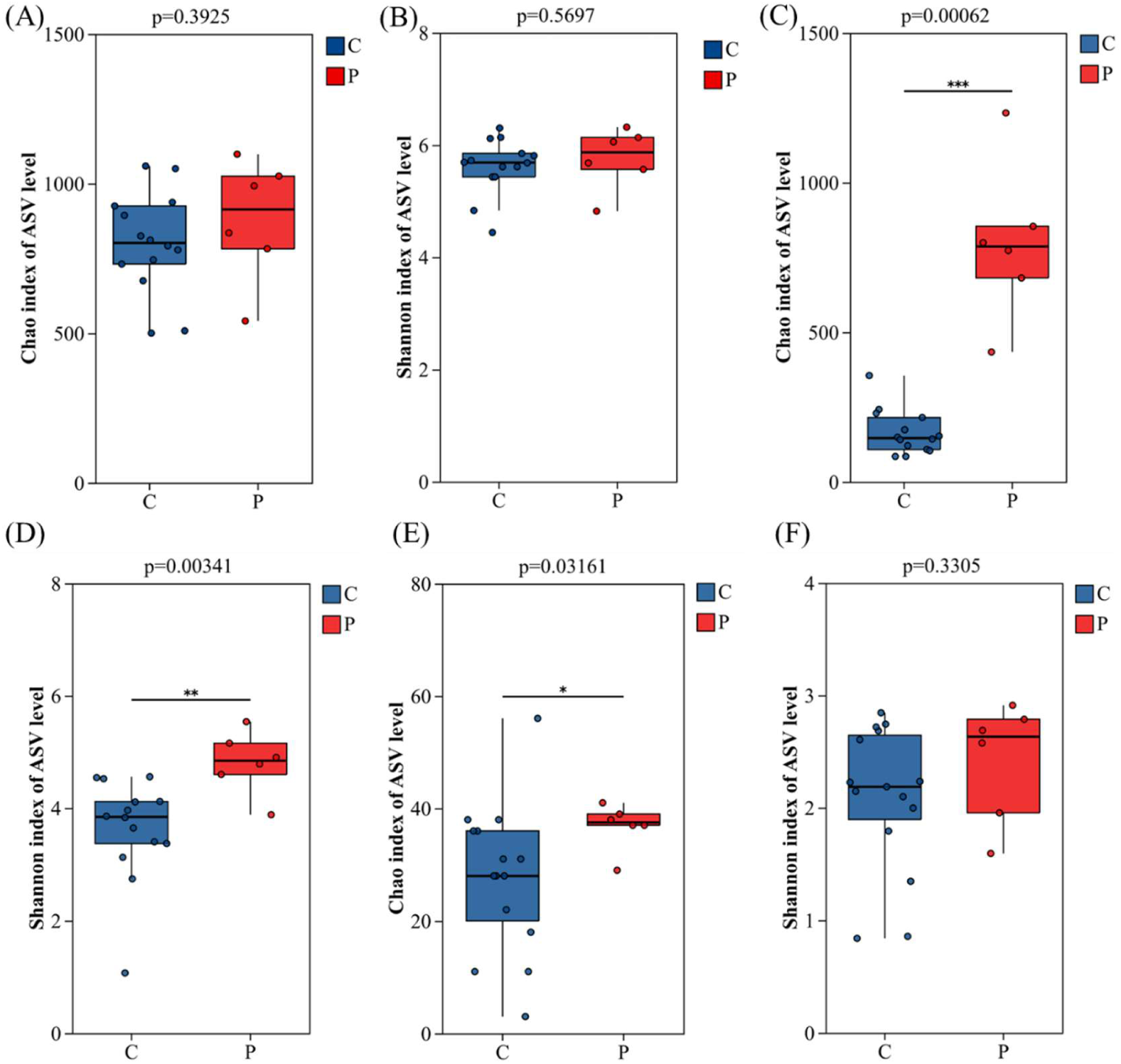
Alpha diversity indices (Chao1 richness and Shannon diversity) of bacterial (A, B), fungal (C, D) and algal (E, F) communities on coastal concrete surfaces (C, *n* = 16) and drainage concrete pipes (P, *n* = 6). Boxes represent the interquartile range (IQR) with the median line; whiskers extend to 1.5 × IQR. Statistical significance was determined by Wilcoxon rank-sum test with false discovery rate (FDR) correction. \**p* < 0.05; \*\**p* < 0.01.

PCoA based on Bray-Curtis distances and ANOSIM (*P* < 0.05) test revealed significant differences in bacterial and algae communities between coastal concrete (C) and concrete pipes (P), with a *p*-value of 0.001 and 0.007, respectively, while no significant differences was observed for fungal communities (Figure 2A-C). This indicates that the compositional differentiation of bacteria and algae may be drive by the environmental factors, whereas fungal composition is relatively uniform across the two habitats.

**Fig. 2.**
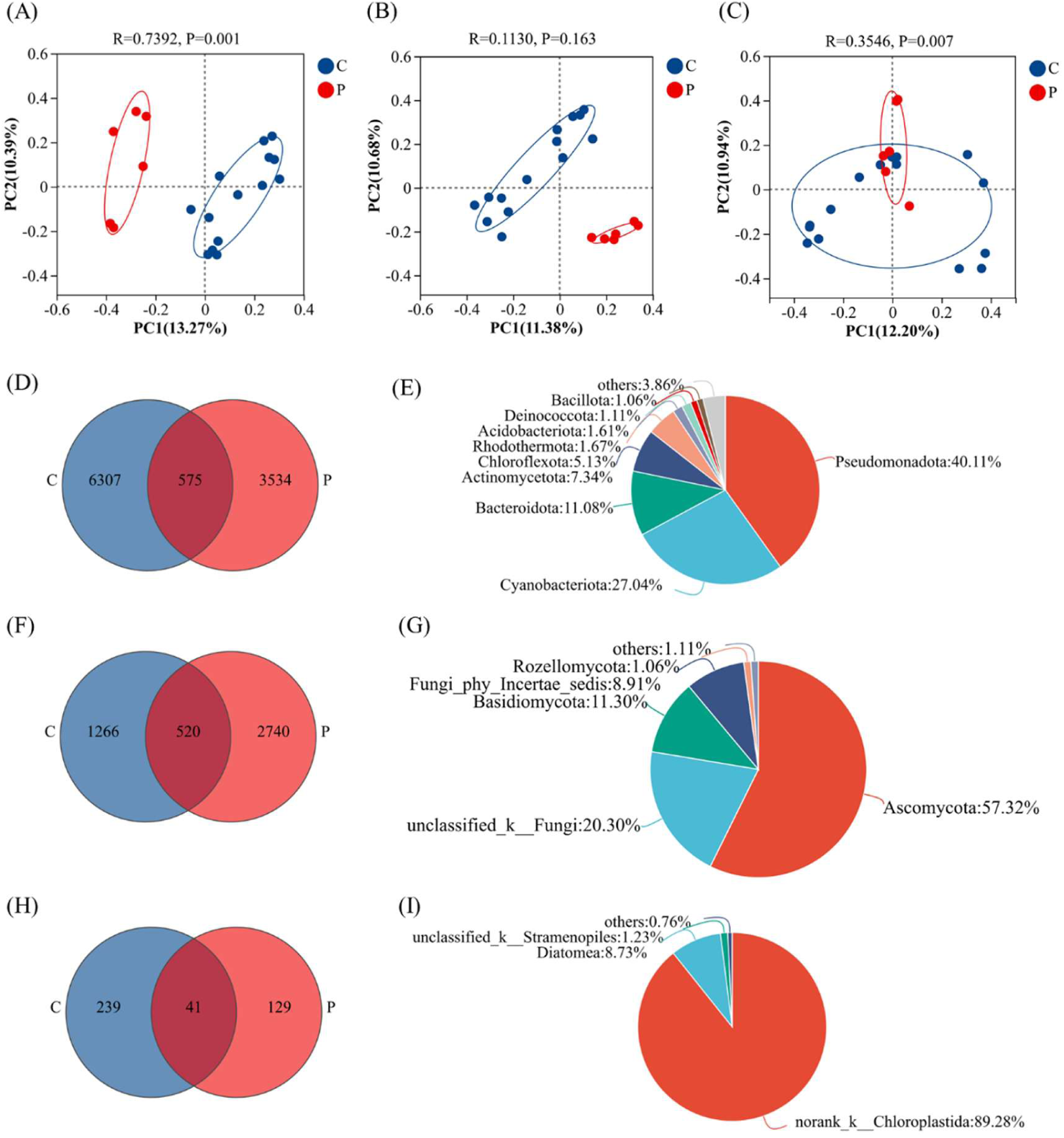
Beta diversity and shared ASVs between coastal concrete (C) and concrete pipes (P). (A–C) Principal coordinate analysis (PCoA) plots based on Bray-Curtis distances for bacteria (A), fungi (B) and algae (C). PERMANOVA p-values (999 permutations) are shown. (D–E) Venn diagram showing shared and unique ASVs for bacteria (D), with the five most abundant phyla of the core ASVs listed (E). (F–G) Same for fungi, and (H–I) for algae.

To compare the microbial compositions of C and P, Venn diagrams were constructed. A total of 575 bacterial ASVs were shared between C and P, most belong to Pseudomonadota (40.11%), Cyanobacteriota (27.04%), Bacteroidota (11.08%), Actinomycetota (7.34%), and Chloroflexota (5.13%)(Figure 2D-E); A total of 520 fungi ASVs were shared, most belong to the phal of Ascomycota (57.32%) and Basidiomycota (11.30%) (Figure 2F-G). A total of 40 algal ASVs were shared, most belong to Chloroplastida (89.26%) and Diatomea (8.75%), respectively (Figure 2H-I). These core members that span across different substrates may constitute the ubiquitous microbial “seed bank” in coastal concrete environments.

### 3.2 Microbial compositions on different concrete surfaces

#### 3.2.1 Bacteria

At the phylum level (Figure 3A), the bacterial communities on both coastal concrete (C) and concrete pipes (P) were dominated by Pseudomonadota, Cyanobacteriota, Bacteroidota, and Actinomycetota, which collectively accounted for over 80% of the relative abundance in C and nearly 80% in P. Notably, Rhodothermota was uniquely enriched in C, whereas Chloroflexota, Acidobacteriota, Deinococcota, Planctomycetota, Nitrospirota, and Bdellovibrionota occurred exclusively in P, all with relative abundance exceeding 0.1% of each sample. Pseudomonadota was the most abundant phylum in both substrates, with higher average relative abundance in C (42.00%) than in P (35.43%), while Cyanobacteriota showed the opposite trend (C: 25.24%; P: 31.06%).

**Fig. 3.**
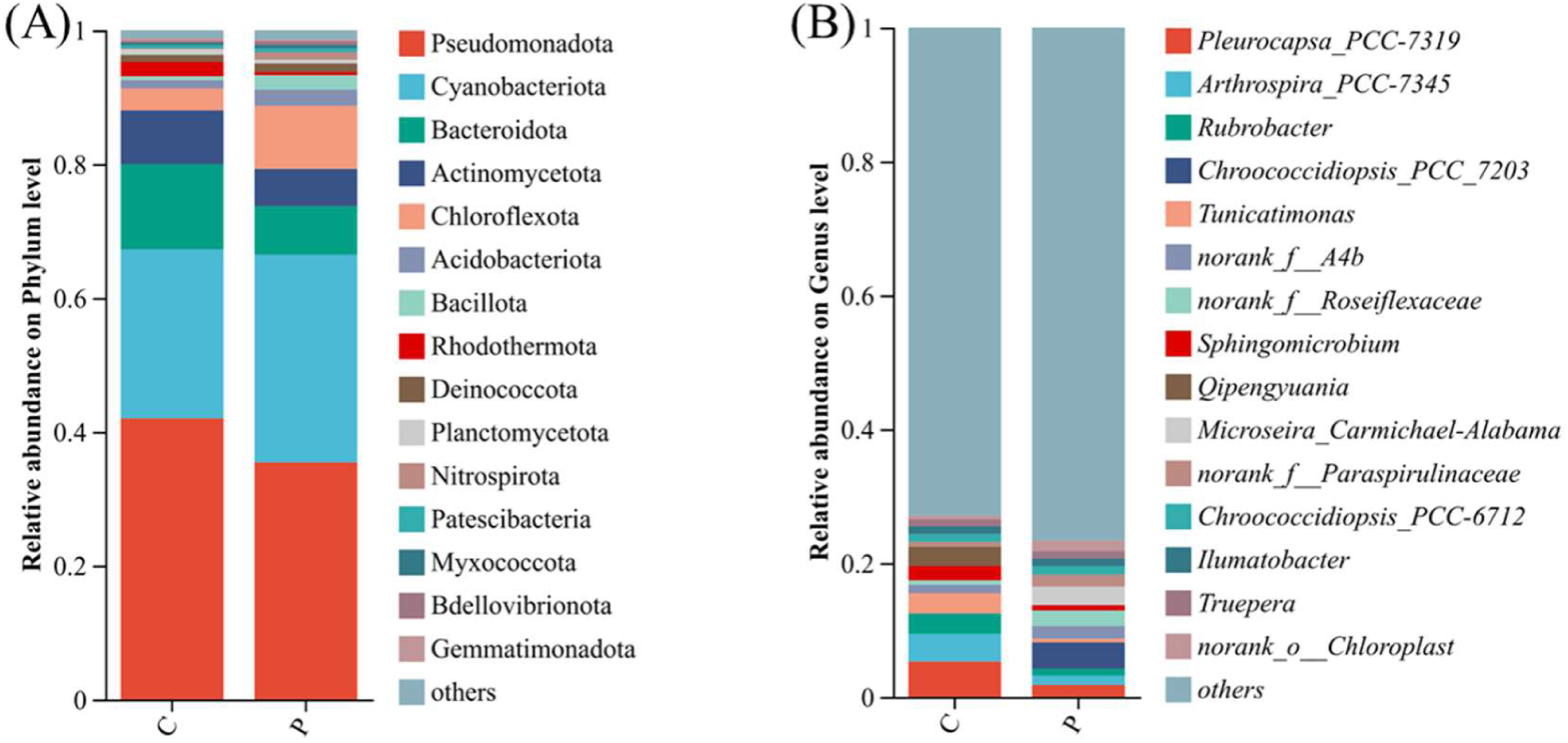
Bacterial community composition at phylum and genus levels. (A) Relative abundance of dominant bacterial phyla (top 10) in C and P. (B) The top 15 most abundant bacterial genera in C and P.

At the genus level (Figure 3B), the overall bacterial community was highly diverse, with over 70% of the relative abundance in both C and P consisted of unclassified taxa or rare genera (<0.1% each). Among the identifiable genera, *unclassified_f Paracoccaceae* (C:9.13%, P:5.26%), *unclassified_f Sphingomonadaceae* (C:9.13%, P:7.68%), *Pleurocapsa_PCC-7319* (C:5.28%, P:1.80%), *Arthrospira_PCC-7345* (C:4.14%, P:1.39%), *Chroococcidiopsis_PCC-6712* (C:1.17%, P:1.26%), and *Ilumatobacter* (C:1.15%, P:1.18%) were shared dominants. *Rubrobacter* (3.05%), *Tunicatimonas* (3.01%), *Sphingomicrobium* (2.17%), and *Qipengyuania* (2.84%) were specifically enriched in C, whereas *Chroococcidiopsis_PCC_7203* (3.96%) was characteristic of P. Despite their dominance, even the most abundant genera (*Pleurocapsa PCC-7319* in C and *Chroococcidiopsis PCC* 7203 in P) had average relative abundances of only 5.28% and 3.96%, respectively, underscoring the high evenness of the bacterial community.

#### 3.2.2 Fungi

The fungal communities were dominated by Ascomycota and Basidiomycota in both substrates, with Ascomycota alone contributing over 50% (C: 58.38%; P: 54.83%), followed was Basidiomycota (C: 14.46%, P: 3.90%). Rozellomycota was an additional dominant phylum uniquely detected in P (1.61%) (Figure 4A).

**Fig. 4.**
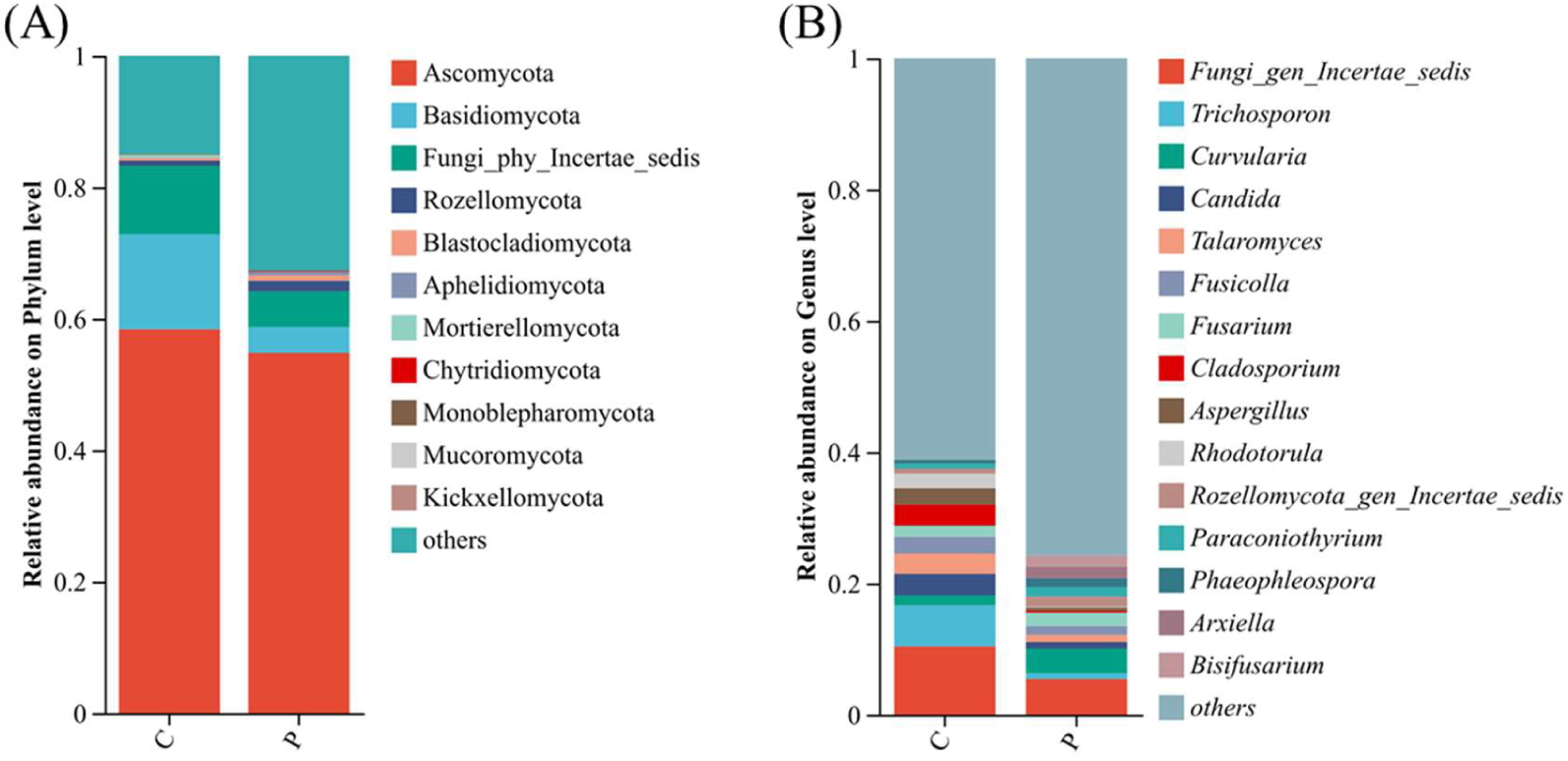
Fungal community composition at phylum and genus levels. (A) Relative abundance of dominant fungal phyla (top 8) in C and P. (B) The top 15 most abundant fungal genera.

At the genus level (Figure 4B), the top 15 dominant genera together accounted for only 38.84% (C) and 24.23% (P) of the relative abundance, indicating a highly diverse fungal community with a long tail of rare taxa. Shared dominants between C and P included *Trichosporon* (C: 6.29%, P:0.89%), *Curvularia* (C: 1.48%, P:3.76%), *Candida* (C: 3.29%, P:0.98%), *Talaromyces* (C: 3.08%, P:1.09%), *Fusicolla* (C: 1.69%, P:2.06%), *Fusarium*(C: 1.69%, P:2.06%), *Paraconiothyrium* (C: 0.78%, P:1.45%), and *Phaeophleospora* (C: 0.56%, P:1.31%). And *Rhodotorula* (2.21%) *Cladosporium* (3.21%), and *Aspergillus* (2.51%) were specifically enriched in C, while *Arxiella* (1.78%) and *Bisifusarium* (1.69%) were specifically enriched in P. As showed above the relative abundance of the dominant genus are all relatively low, indicating that the ecosystem function of the fungal communities is relatively stable and may have a strong ability to resistance to disturbance.

#### 3.2.3 Algae

The algal communities were overwhelmingly dominated by the phylum norank_k Chloroplastida (green algae), which accounted for approximately 90% of the relative abundance in both C (88.33%) and P (91.22%). Diatoms (Ochrophyta) was a secondary phylum in C (10.66%) but negligible in P (Figure 5A).

**Fig. 5.**
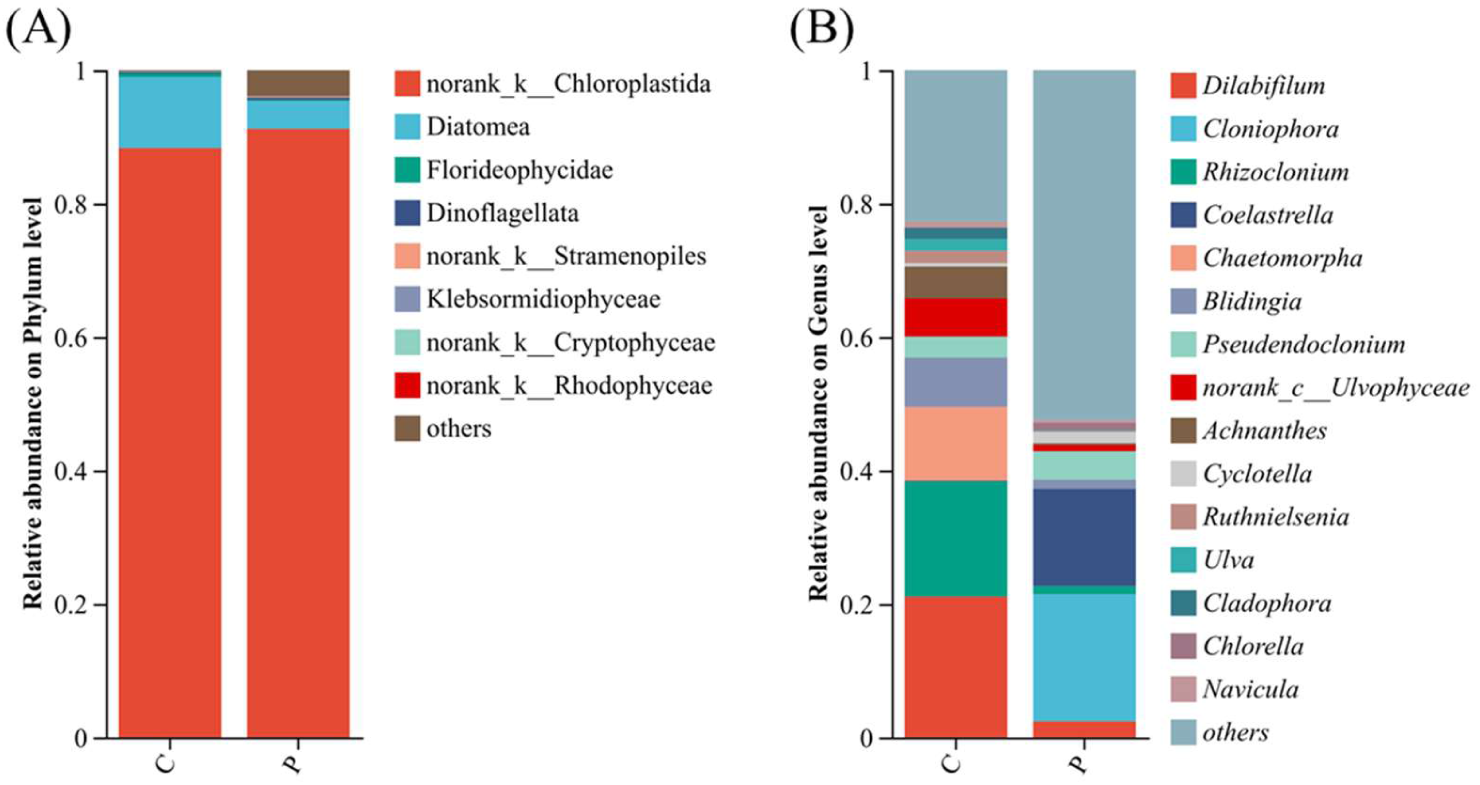
Algal community composition at phylum and genus levels. (A) Relative abundance of algal phyla in C and P. (B) The top 15 most abundant algal genera.

At the genus level (Figure 5B), *Dilabifilum* (C:21.11%, P:2.42%) and *Pseudendoclonium* (C:3.14%, P:4.26%), and *Pseudendoclonium* (C:3.14%, P:4.26%) were shared dominants. Coastal concrete surfaces (C) were characterized by the enrichment of *Chaetomorpha* (11.05%), *Blidingia* (7.39%), *Pseudendoclonium* (3.14%), *Achnanthes*(4.79%), *Ruthnielsenia*(1.87%), and *Ulva*(1.73%), with *Dilabifilum* (21.11%) and *Rhizoclonium* (17.25%) being the most abundant. In contrast, pipe concrete surfaces (P) were dominated by *Cloniophora* (19.06%), *Coelastrella* (14.58%), *Cyclotella* (1.69%) and *Chlorella* (1.20%). The scarcity of shared dominant genera and the prevalence of unique dominants indicate strong environmental filtering and niche differentiation driven by these two distinct habitat conditions for algae.

### 3.3 Microbial differences between C and P

To determine the microbial that exhibited significant differences between the concrete and concrete pipes, Student’s *t*-test was conducted between the two groups at genus level. The results showed that the relative abundances of *Chroococcidiopsis_PCC_7203*, *Microseira_Carmichael-Alabama*, *norank_o Chloroplast*, *Planktothrix_NIVA-CYA_15*, and bacterial genus *Sphingomonas* were significantly lower in C than in P, whereas *Tunicatimonas*, *Sphingomicrobium*, *Qipengyuania*, *orank_f Rhodothermaceae*, *Muricauda* exhibited the opposite trend (Figure 6A)

**Fig. 6.**
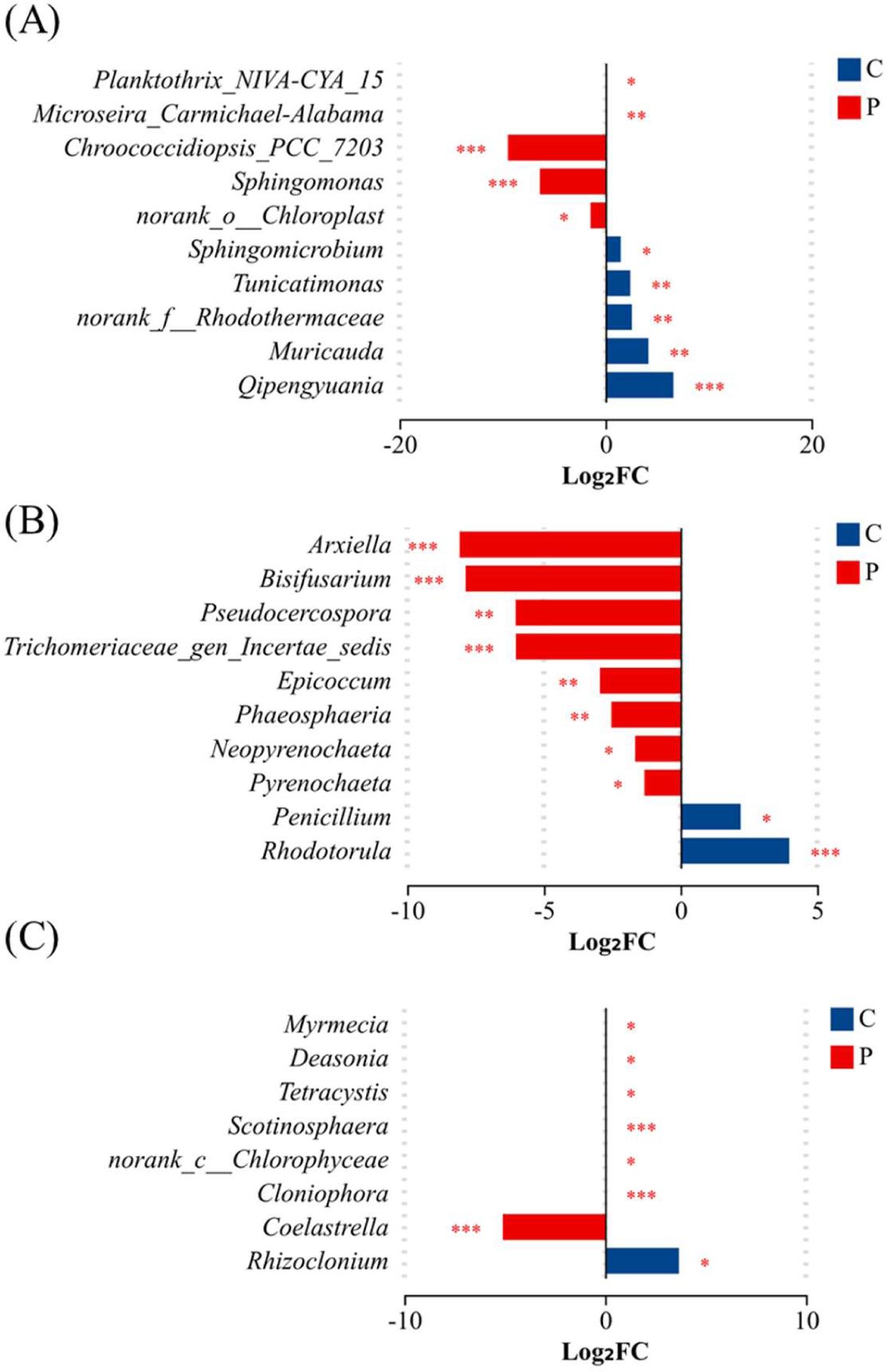
Differential abundant microbial genera between C and P identified by Student’s t-test with FDR correction (q < 0.05). (A) Top 10 bacterial genera with significantly different relative abundances. (B) Top 10 fungal genera. (C) Top 10 algal genera. Bars represent mean relative abundance ± standard deviation. Blue bars indicate enrichment in C, red bars indicate enrichment in P.

For fungi, the relative abundances of most top 10 like *Arxiella, Bisifusarium*, *Trichomeriaceae_gen_Incertae_sedis*, *Phaeosphaeria*, *Neopyrenochaeta*, *Pseudocercospora*, *Pyrenochaeta* were significantly lower in C than in P, and only *Fusicolla*, *Rhodotorula*, and *Penicillium* showed higher relative abundances in C than in P (Figure 6B)

As for algae, the relative abundances of *Coelastrella* were significantly higher in C than in P, while the other algae genus of the top 10 like *Myrmecia*, *Deasonia*, *Tetracystis*, *Scotinosphaera*, *norank_c Chlorophyceae*, *Cloniophora*, *Rhizoclonium* were significantly lower in C than in P. (Figure 6C)

These differential patterns indicate that the concrete pipes environment (P), characterized by continuous water flow, persistent humidity, and organic nutrient accumulation, favors oligotrophic degraders like *Sphingomonas*, saprotrophic and acid-producing fungi like *Arxiella* and *Bisifusarium*, as well as stress-sensitive green algae like *Myrmecia* and *Cloniophora*. In contrast, the costal concrete surface (C), subject to periodic desiccation and salt stress, selects for halotolerant EPS-producing bacteria like *Tunicatimonas* and *Muricauda*, a few moisture-independent fungi like *Penicillium* and *Rhodotorula*, and the extremotolerant alga *Coelastrella*, whose desiccation and UV resistance enable its unique enrichment on exposed concrete.

### 3.4 The environmental factors and db-RDA around C and P

The water physicochemical parameters of the two habitats are show in figure 7A-F. The pH of the coastal concrete water (C) was 7.93, slightly higher than that of the concrete pipes water (P) at 7.80. Salinity in C (32.34‰) was substantially higher than in P (9.55‰), reflecting the marine influence. Temperature was comparable between the two sites (34.49 °C in C vs. 35.18 °C in P). In contrast, concentrations of NH_4_^+^-N, NO_3_^-^ -N and chemical oxygen demand (COD) were markedly higher in P (0.46 mg/L, 0.76 mg/L and 2.40 mg/L, respectively) than in C (0.02 mg/L, 0.22 mg/L and 0.55 mg/L, respectively), indicating that the pipe environment carried a higher organic and nutrient load. These distinct physicochemical profiles likely exert strong selective pressures on the microbial communities colonizing the concrete surfaces.

**Fig. 7.**
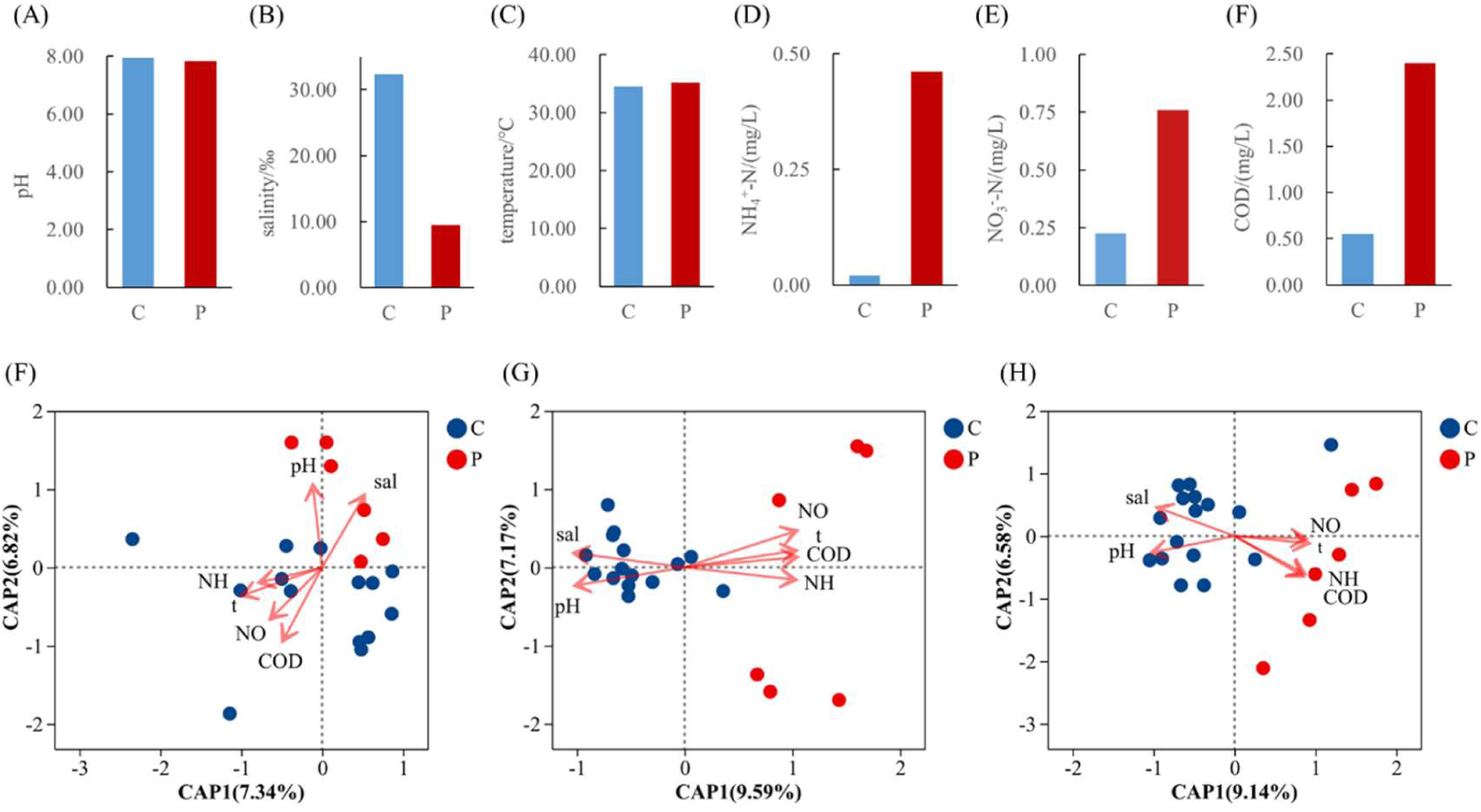
Water physicochemical parameters and distance-based redundancy analysis (db-RDA) with sampling sites. (A–F) Comparison of pH (A), salinity (B), temperature (C), NH_4_^+^-N (D), NO_3_^-^-N (E) and chemical oxygen demand (COD, F) between C and P. Boxplot details as in Fig. 1. (G–I) db-RDA ordination plots for bacteria (G), fungi (H) and algae (I) based on Bray-Curtis distances. Significant environmental variables (*p* < 0.05, permutation test) are shown as vectors. The explained variances of db-RDA axes are indicated.

The distance-based redundancy analysis (db-RDA) based on Bray-Curtis distances was used to evaluate the influence of environmental factors on the microbial community composition. For bacteria (Figure 7G), salinity (*p* = 0.030), NH_4_^+^-N (*p* = 0.007), NO_3_^-^-N (*p* = 0.016) and COD (*p* = 0.002) significantly shaped the bacterial community structure, whereas temperature showed a marginally non-significant effect (*p* = 0.066). For fungi (Figure 7H), all tested variables exhibited significant effects: pH (*p* = 0.028), salinity (*p* = 0.001), temperature (*p* = 0.005), NH_4_^+^-N (*p* = 0.001), NO_3_^-^-N (*p* = 0.001) and COD (*p* = 0.001). For algae (Figure 7I), salinity (*p* = 0.001), temperature (*p* = 0.005), NH_4_^+^-N (*p* = 0.001), NO_3_^-^-N (*p* = 0.001) and COD (*p* = 0.001) were significant drivers, while pH was not significant (*p* = 0.23). Under the influence of these variables, the samples clearly separated into two distinct clusters corresponding to coastal concrete (C) and concrete pipes (P). The bacterial community associated with C was mainly driven by NH_4_^+^-N, NO_3_^-^-N, temperature and COD, while the fungal and algal communities in C were predominantly influenced by salinity and pH. In contrast, the P samples exhibited opposite relationships with these environmental gradients, indicating a distinct ecological niche. These results demonstrate that environmental filtering by salinity, nutrient availability and pH differentially structures the bacterial, fungal and algal communities on coastal concrete and concrete pipe surfaces.

### 3.5 Co-occurrence network analysis among different microorganisms

To further explore the potential ecological interactions among microbial taxa in C and P surfaces, we constructed co-occurrence networks based on Spearman’s rank correlation coefficients (|*ρ*| > 0.5, *p* < 0.05), including co-occurrence networks based on the combined of top 15 genera of bacteria, fungi, and algae in each habitat of C and P; as well as the top 45 genera of bacteria, fungi, and algae in C and P, respectively.

The co-occurrence networks of the top 15 genera of bacteria, fungi, and algae in C comprised 33 nodes and 80 edges (Figure 8A), while the network in P consisted of 34 nodes and 89 edges (Figure 8B). The positive correlation ratios measured in C and P were 57.50% and 56.18% respectively, indicating that both interspecies competition and cooperation coexist, with a slight predominance of competition. Keystone taxa identified by high betweenness centrality (BC) included *Achnanthes* (BC = 0.192), *Chaetoceros* (BC = 0.142), *Candida* (BC = 0.135), and *Tunicatimonas* (BC = 0.107) in C, whereas *Amoeboaphelidium* (BC = 0.282), *Cloniophora* (BC = 0.212), *Bisifusarium* (BC = 0.162), *Tunicatimonas* (BC = 0.116), and *Aspergillus* (BC = 0.113) in P, suggesting their pivotal roles in bridging microbial modules in C and P biofilm network, respectively.

**Fig. 8.**
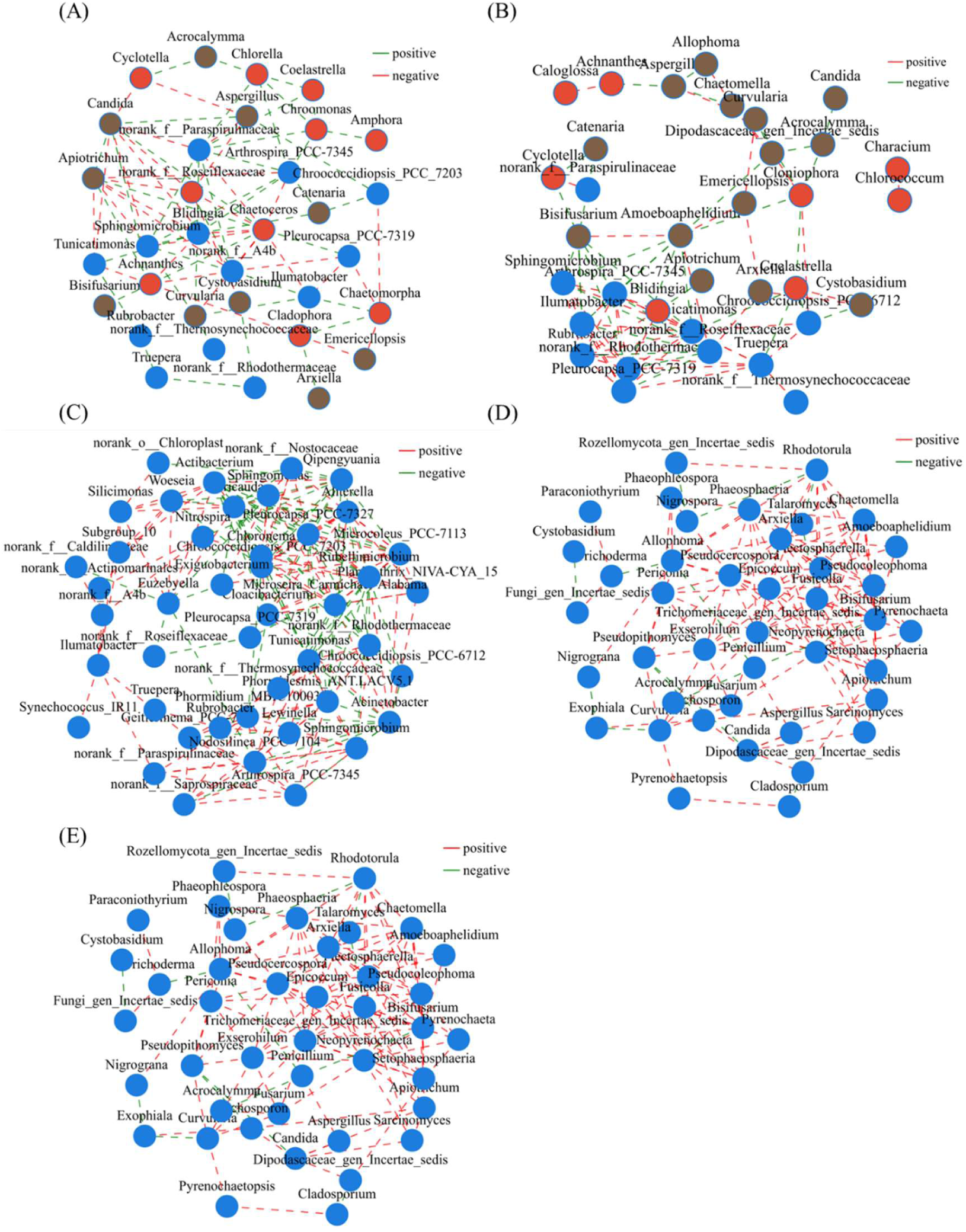
Co-occurrence networks of microbial taxa based on Spearman’s rank correlations (|ρ| > 0.5, p < 0.05). (A) Cross-domain network using the top 15 most abundant genera of bacteria, fungi and algae in C. (B) Same for P. (C) Bacterial network using the top 45 most abundant genera in C and P combined. (D) Fungal network (top 45 genera). (E) Algal network (top 45 genera). Node size is proportional to betweenness centrality; edge color indicates positive (green) or negative (red) correlation.

For the co-occurrence networks of the top 45 bacteria genera in C and P contained of 44 nodes and 199 edges (Figure 8C), while the network of the top 45 fungal genera consisted of 41 nodes and 132 edges (Figure 8D), and the network of the top 45 algal genera in C and P was the most compartmentalized, with 41 nodes but only 74 edges (Figure 8E), reflecting a less integrated algal community compared to bacteria and fungi. The positive co-expression ratios measured of the top 45 genera of bacteria, fungi, and algae in C and P, were 68.81%, 89.39%, and 97.30% respectively, indicating stronger cooperative interactions within the same kingdom, especially among algae. Keystone taxa included bacterial genera *Rubellimicrobium* (BC = 0.106), *Euzebyella* (BC = 0.101), and *Ilumatobacter* (BC = 0.093); fungal genera *Allophoma* (BC = 0.252), *Trichoderma* (BC = 0.145), and *Rhodotorula* (BC = 0.117); and algal genera *Prototheca* (BC = 0.222), *Desmodesmus* (BC = 0.205), *Ulvella* (BC = 0.140), and *Navicula* (BC = 0.091).

### 3.6 Functional prediction of bacterial community on C and P

To gain insight into the potential metabolic capabilities of bacterial communities in C and P, we compared the proportions of KEGG functional pathways using a Wilcoxon rank-sum test. According to the KEGG hierarchy, Level 1 pathways (Figure 9A) are categorized into five functional categories, in which Cellular Processes (*p* = 0.015) and Environmental Information Processing (*p* = 0.012) were significantly higher in P than in C, whereas Metabolism (*p* = 0.013) was significantly higher in C than in P. This indicates that C habitats favor core metabolic activities, while P habitats prioritize cellular regulation and environmental sensing.

**Fig. 9.**
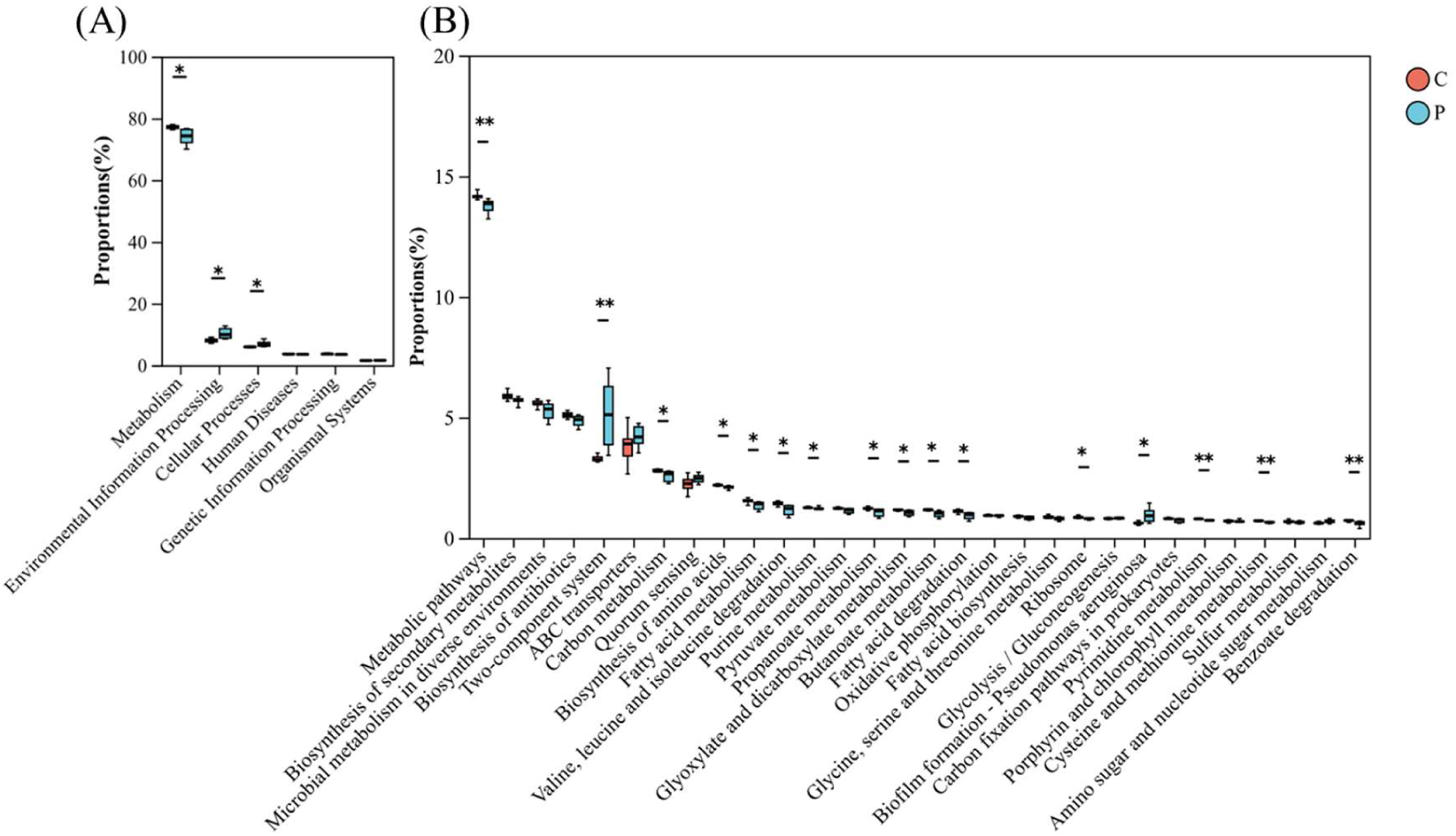
Functional predictions of bacterial communities using PICRUSt2. (A) KEGG Level 1 pathways: relative abundance of five functional categories in C and P. (B) Top 30 KEGG Level 3 pathways between C and P. Error bars represent standard deviation. \**p* < 0.05; \*\**p* < 0.01.

Nearly half of the Level 3 pathways (Figure 9B) were significantly higher in C than in P, and most of this belonged to Metabolism Level 1 pathways, such as metabolic pathways (*p* = 0.001), carbon metabolism (*p* = 0.019), biosynthesis of amino acids (*p* = 0.003), fatty acid metabolism (*p* = 0.006), valine, leucine and isoleucine degradation (*p* = 0.003), propanoate metabolism (*p* = 0.004), glyoxylate and dicarboxylate metabolism (*p* = 0.007), butanoate metabolism (*p* = 0.003), fatty acid degradation (*p* = 0.003), cysteine and methionine metabolism (*p* = 0.002), and benzoate degradation (*p* = 0.002). In contrast, pathways like Two-component system (*p* = 0.001) and Biofilm formation - Pseudomonas aeruginosa (*p* = 0.009) under level 1 pathways of Environmental Information Processing and Cellular Processes, respectively, were significantly lower in C than P. Notably, sulfur metabolism did not differ significantly (*p* = 0.174), possibly due to the low overall abundance of sulfur-cycling taxa at the community level, despite their presence in the taxonomic analysis (Section 3.4.1). All of this indicated that bacterial community in C emphasizes active metabolic pathways, including degradative and biosynthetic, while P prioritizes regulatory and biofilm-forming strategies to exert the functional roles.

### 3.7 Microbial taxa potentially involved in concrete deterioration and healing

To identify potential microbial taxa associated with microbiologically influenced concrete corrosion (MICC) (based on relevant research literature, Supplementary), we classified bacteria, fungi, and algae based on their relative abundance and occurrence in C and P samples (Tables 1-2). The classification distinguished between abundant (≥ 1% relative abundance in at least one sample) and permanently rare (always ≤1% relative abundance) taxa, further categorized by distribution breadth: broad (≥75% occurrence), intermediate (>10% and <75% occurrence), and narrow (≤ 10% occurrence)(34).

**Table 1.** Microorganisms potentially involved in concrete deterioration.

| Categories |  | Genus |  |
| --- | --- | --- | --- |
| Primary categories | Secondary categories | C | P |
| Abundant | Broad | B: *Rubrobacter*<br>F : *Cladosporium*,<br>*Aspergillus*,<br>*Penicillium*,<br>*Trichoderma*<br>A: *Rhizoclonium* | B: -<br>F: *Epicoccum*, *Aspergillus*,<br>*Fusarium*<br>A: - |
|  | Intermediate | B: *Vibrio*, *Halomonas*,<br>*Sphingomonas*<br>F: *Lasiodiplodia*,<br>*Epicoccum*, *Alternaria*, | B: *Pseudomonas*<br>F: -<br>A: *Rhizoclonium* |
|  |  | *Exophiala*,<br>*Acremonium*,<br>*Fusarium*,<br>*Chaetomium*,<br>*Rhodotorula*<br>A: *Chaetomorpha*,<br>*Cladophora*, *Ulva*,<br>*Achnanthes*, *Navicula* |  |
|  | Narrow | B: -<br>F: *Purpureocillium*<br>A: - | B: *Paenibacillus*<br>F: -<br>A: - |
| Permanently<br>rare | Broad | B: *Nitrospira*,<br>*Clostridium*<br>F: -<br>A: - | B: -<br>F: *Lasiodiplodia*,<br>*Cladosporium*,<br>*Aureobasidium*, *Alternaria*,<br>*Exophiala*, *Penicillium*,<br>*Trichoderma*, *Acremonium*,<br>*Rhodotorula*<br>A: *Chlorella* |
|  | Intermediate | B: *Mycobacterium*,<br>*Pseudomonas*,<br>*Sulfurovum*,<br>*Nitrosomonas*,<br>*Arcobacter*,<br>*Corynebacterium*,<br>*Methylobacterium*<br>F: *Aureobasidium*,<br>*Serpula*<br>A: *Chlorella*, *Nitzschia* | B: *Rubrobacter*,<br>*Halomonas*, *Sphingomonas*,<br>*Mycobacterium*,<br>*Clostridium*, *Sulfurovum*,<br>*Staphylococcus*, *Thiothrix*,<br>*Streptomyces*, *Alkaliphilus*,<br>*Nocardia*, *Enterococcus*,<br>*Thiobacillus*<br>F: *Phoma*, *Lecanicillium*,<br>*Purpureocillium*, |
|  |  |  | *Stachybotrys*, *Chaetomium*,<br>*Serpula*<br>A: *Klebsormidium*, *Ulva*,<br>*Achnanthes*, *Navicula*,<br>*Nitzschia*, *Melosira* |
|  | Narrow | B: *Staphylococcus*,<br>*Thiothrix*,<br>*Desulfovibrio*,<br>*Streptomyces*,<br>*Paenibacillus*,<br>*Massilia*, *Alkaliphilus*,<br>*Desulfobulbus*,<br>*Nocardia*<br>F: *Stachybotrys*<br>A: - | B: -<br>F: -<br>A: - |

Abundant: $\geq 1\%$ relative abundance in at least one sample. Permanently rare: relative abundance always $\leq 1\%$ . Broad: $\geq 75\%$ occurrence, Intermediate: $>10\%$ and $<75\%$ samples, and Narrow: $\leq 1\%$ samples. B: Bacteria, F: fungi, A: Algae.

**Table 2.** Microorganisms that may be involved in concrete healing.

| Categories |  | Genus |  |
| --- | --- | --- | --- |
| Primary categories | Secondary categories | C | P |
| Abundant | Broad | B: *Exiguobacterium* | B: - |
|  |  | F: *Aspergillus*,<br>*Penicillium*, *Trichoderma* | F: *Aspergillus*, *Candida*,<br>*Fusarium* |
|  | Intermediate | B: *Halomonas*,<br>*Pseudoalteromonas*<br>F: *Candida*, *Fusarium*,<br>*Rhodotorula* | B: *Pseudomonas*<br>F: - |
| Permanently<br>rare | Broad | B: -<br>F: - | B: *Pseudoalteromonas*,<br>*Sutcliffiella*,<br>*Paenibacillus*<br>F: *Trichoderma*,<br>*Penicillium*, *Rhodotorula* |
|  | Intermediate | B: *Bacillus*,<br>*Pseudomonas*,<br>*Paracoccus*,<br>*Corynebacterium*,<br>*Arthrobacter*<br>F: - | B: *Exiguobacterium*,<br>*Halomonas*, *Bacillus*,<br>*Halobacillus*,<br>*Rhodococcus*,<br>*Micrococcus*,<br>*Bhargavaea*,<br>*Staphylococcus*,<br>*Arthrobacter*,<br>*Chryseomicrobium*,<br>*Streptomyces*,<br>*Sporosarcina*,<br>*Anaerobacillus*,<br>*Enterococcus*<br>F: *Neurospora*,<br>*Naganishia* |
|  | Narrow | B: *Halobacillus*,<br>*Rhodococcus*,<br>*Sutcliffiella*, *Micrococcus*, | B: -<br>F: - |
|  |  | *Bhargavaea*,<br>*Staphylococcus*,<br>*Chryseomicrobium*,<br>*Streptomyces*,<br>*Paenibacillus*,<br>*Sporosarcina*<br>F: - |  |

Abundant: $\geq 1\%$ relative abundance in at least one sample. Permanently rare: relative abundance always $\leq 1\%$ . Broad: $\geq 75\%$ occurrence, Intermediate: $>10\%$ and $<75\%$ samples, and Narrow: $\leq 1\%$ samples. B: Bacteria, F: fungi, A: Algae.

#### 3.7.1 Microbial taxa potentially involved in concrete deterioration

Based on literature reports, we curated a list of taxa from our sequencing data that have been previously implicated in microbially induced concrete corrosion (MICC). Their occurrence frequencies and relative abundances were stratified into abundant (≥1% in at least one sample) versus permanently rare (always ≤1%), and further categorized by spatial occupancy (broad: ≥75% samples; intermediate: >10% to <75%; narrow: ≤1% samples) (Table 1).

For abundant bacteria, *Rubrobacter* was the only broad-occurrence taxon potentially linked to MICC, and it was found exclusively on coastal concrete surfaces (C). In contrast, concrete pipe surfaces (P) harbored no abundant broad-distribution bacteria but contain *Pseudomonas* as an intermediate-occurrence taxon. *Vibrio*, *Halomonas*, and *Sphingomonas* were also identified as intermediate occurrence bacteria in C. Among permanently rare bacteria, C supported a narrow-distribution guild, such as *Staphylococcus*, *Thiothrix*, *Desulfovibrio*, *Streptomyces*, *Paenibacillus*, *Massilia*, *Alkaliphilus*, *Desulfobulbus*, and *Nocardia*, whereas P lacked narrow-distribution taxa but hosted a diverse set of intermediate-distribution rare bacteria, including *Rubrobacter*, *Halomonas*, *Sphingomonas*, *Clostridium*, *Staphylococcus*, *Thiothrix*, *Streptomyces*, *Alkaliphilus*, *Nocardia*, *Enterococcus*, and *Thiobacillus*. This pattern suggests that MICC-related bacteria are narrower niche in C, while adopt broader environmental tolerances in P, possibly reflecting the more stable, nutrient-rich conditions inside pipes.

For abundant fungi, C exhibited a broader suite of broad-distribution taxa, including *Cladosporium*, *Aspergillus*, *Penicillium*, and *Trichoderma*, compared to P, which contained *Epicoccum*, *Aspergillus*, and *Fusarium*. Additionally, C contained numerous abundant intermediate-distribution fungi, such as *Lasiodiplodia*, *Epicoccum*, *Alternaria*, *Chaetomium*, *Rhodotorula*, whereas P had none in this category.Among permanently rare fungi, the opposite was true: P harbored a substantial broad-distribution rare fungal community, such as *Lasiodiplodia*, *Cladosporium*, *Aureobasidium*, *Alternaria*, *Exophiala*, *Penicillium*, *Trichoderma*, *Acremonium*, and *Rhodotorula*, while C had none. P also hosted a richer assemblage of intermediate-distribution rare fungi, including *Phoma*, *Lecanicillium*, *purpureocillium*, *Stachybotrys*, *Chaetomium*, and *Serpula*, while C only had two of these taxa, including *Aureobasidium* and *Serpula*. These findings indicate that pipe concrete environments maintain a larger reservoir of rare but widely distributed fungi with documented corrosive potential (e.g., organic acid producers), whereas coastal concrete surfaces favor abundant, corrosion-related fungi that are more consistently present.

For abundant algae involved in MICC were almost exclusively associated with C: *Rhizoclonium* as a broad-distribution taxon, and *Chaetomorpha*, *Cladophora*, *Ulva*, *Achnanthes*, and *Navicula* as intermediate-distribution taxa. P only contained *Rhizoclonium* as an intermediate-distribution abundant alga. In the permanently rare algal pool, P uniquely harbored the broad-distribution taxon *Chlorella* and a diverse set of intermediate-distribution rare algae, including *Klebsormidium*, *Ulva*, *Achnanthes*, *Navicula*, *Nitzschia*, and *Melosira*). C had only *Chlorella* and *Nitzschia* as intermediate-distribution rare algae. The higher diversity and wider occurrence of rare MICC-related algae in P suggest that pipes provide more favorable hydration and light regimes for less competitive phototrophs, which may contribute to biofilm-mediated corrosion through oxygen concentration cells and localized pH changes.

#### 3.7.2 Microbial taxa potentially involved in concrete healing

In addition to corrosion-related taxa, we identified a range of microorganisms with documented potential for concrete healing, primarily through microbially induced calcium carbonate precipitation (MICP) or biofilm-mediated crack sealing (Table 2). For bacteria, among abundant taxa, *Exiguobacterium* was the only broad-distribution healing-associated bacterium, and it occurred exclusively on coastal concrete surfaces (C). C also harbored *Halomonas* and *Pseudoalteromonas* as abundant intermediate-distribution taxa, whereas P contained only *Pseudomonas* as intermediate-distribution in abundant category. In the permanently rare pool, P hosted three broad-distribution rare healing bacteria, including *Pseudoalteromonas*, *Sutcliffiella*, and *Paenibacillus*, while C had none. Moreover, P exhibited a highly diverse assemblage of intermediate-distribution rare healing bacteria such as *Exiguobacterium*, *Halomonas*, *Bacillus*, *Halobacillus*, *Rhodococcus, Micrococcus*, *Bhargavaea*, *Staphylococcus*, *Arthrobacter*, *Chryseomicrobium*, *Streptomyces*, *Sporosarcina*, *Anaerobacillus*, and *Enterococcus*, far outnumbering those in C which only contained *Bacillus*, *Pseudomonas*, *Paracoccus*, *Corynebacterium*, and *Arthrobacter*. Conversely, C uniquely retained a narrow-distribution rare guild, including *Halobacillus*, *Rhodococcus*, *Sutcliffiella*, *Micrococcus*, *Bhargavaea*, *Staphylococcus*, *Chryseomicrobium*, *Streptomyces*, *Paenibacillus*, and *Sporosarcina*, which was absent in P. This pattern indicates that concrete surfaces (C) contain a few abundant and highly specialized (narrow-distribution) rare healing bacteria, whereas concrete pipes (P) lack abundant healers but maintain a large and broadly distributed reservoir of rare healing taxa.

For abundant fungi, C and P shared *Aspergillus* as a broad-distribution healing taxon. Additionally, C contained *Penicillium* and *Trichoderma* as broad-distribution taxa, while P uniquely harbored *Candida* and *Fusarium* in this category. C also possessed abundant intermediate-distribution healing fungi include *Candida*, *Fusarium*, and *Rhodotorula*, whereas P had none. Among permanently rare fungi, P exclusively harbored three broad-distribution taxa include *Trichoderma*, *Penicillium*, *Rhodotorula* and two intermediate-distribution taxa include *Neurospora* and *Naganishia*; C showed no rare healing fungi. Thus, fungal healing potential is consistently present as abundant taxa on concrete surfaces, whereas in pipes it is relegated to a rare but widely distributed reservoir.

## 4 Discussion

### 4.1 The composition of microbial communities is selected by the environment, resulting in different functions and ecological mechanisms

#### 4.1.1 Microbial diversity and dominant taxa are shaped by environmental conditions

The α-diversity analyses revealed that fungal and algal communities in P exhibited significantly higher richness (Chao1) and diversity (Shannon) than those in C, whereas bacterial diversity did not differ significantly between the two substrates. This differential response suggests that fungi and algae are more sensitive to substrate type and associated environmental conditions, while bacteria may possess stronger cross-matrix adaptability, likely due to their physiological plasticity and rapid colonization capabilities (8). The pipe environment, characterized by continuous water flow, persistent high humidity, and elevated organic nutrient loads (NH_4_^+^-N, NO_3_^-^-N, and COD), provides stable and resource-rich niches that support a wider range of saprotrophic fungi and diverse green algae (10, 35). In contrast, coastal concrete surfaces experience periodic desiccation, high salinity, and solar radiation, which restrict colonization to extremotolerant taxa(1, 36).

Despite the marked environmental differences between C and P, a large core of ASVs was shared between the two substrates, **including** 575 bacterial ASVs, 520 fungal ASVs and 40 algal ASVs, which may constitute a ubiquitous microbial “seed bank” in both habitats. These core members, belonged predominantly to a few shared phyla due to their high phenotypic plasticity and environmental tolerance (8, 37), may maintain essential functions such as primary colonization and nutrient cycling across different concrete infrastructures (2, 3, 6, 38). Furthermore, PCoA and ANOSIM analyses revealed that the community compositions of bacteria and algae were significantly separated between C and P (*p* = 0.001 and *p* = 0.007, respectively), while fungal composition was relatively uniform. This implies that environmental filtering exerts a stronger effect on bacteria and algae; fungi, owing to their efficient spore dispersal mechanisms and broad ecological amplitudes, may exhibit a more homogeneous distribution across the two substrates (10).

Student’s *t*-test at the genus level identified three consistent ecological patterns. For bacteria, oligotrophic degraders like *Sphingomonas* were significantly more abundant in P, consistent with the nutrient-rich pipe environment (39, 40); conversely, halotolerant EPS-producing bacteria like *Tunicatimonas* and *Muricauda* were enriched in C, reflecting adaptation to high salinity and desiccation stress (9, 41). For fungi, saprotrophic and acid-producing genera like *Arxiella* and *Bisifusarium* dominated in P, aligning with the organic-rich, slightly acidic sewer conditions (10, 21), whereas *Penicillium*, which is known for its tolerance low water activity and UV radiation, was enriched in C (11, 42).For algae, the extremotolerant green alga *Coelastrella* that desiccation- and UV-resistant was characteristic of C, while most other differentially abundant algae like *Myrmecia* and *Cloniophora* were more prevalent in P (1, 36). These patterns demonstrate how local environmental conditions-particularly salinity, nutrient availability, and moisture regime-select for distinct microbial functional guilds on each concrete substrate.

The db-RDA analysis explicitly attributed the community differentiation to salinity, NH_4_^+^-N, NO_3_^-^-N, and COD. Salinity was the strongest discriminating factor between C (32.34‰) and P (9.55‰). The high-salinity environment favored the enrichment of halotolerant EPS-producing bacteria like *Tunicatimonas* and *Muricauda* in C (9, 41) and promoted the metabolic activity of sulfur-oxidising and sulfate-reducing bacteria, thereby accelerating sulfur-driven corrosion (4, 37). In contrast, the significantly higher nutrient concentrations of NH_4_^+^-N, NO_3_^-^-N, and COD in pipes supported the growth of oligotrophic degraders like *Sphingomonas* and saprotrophic/acid-producing fungi like *Arxiella* and *Bisifusarium* (10, 39, 40). pH significantly affected only the fungal community (*p* = 0.028), which may be related to the slightly more alkaline environment of coastal concrete of pH 7.93 vs. 7.80 favouring alkaliphilic fungi such as *Aspergillus* and *Penicillium* (11, 42); bacteria and algae did not respond significantly to pH, indicating a broad tolerance range under the investigated conditions.

#### 4.1.2 Functional predictions and co-occurrence networks reveal distinct metabolic and ecological strategies between the two habitats

Beyond compositional differences, KEGG functional prediction of bacterial communities revealed that metabolism-related pathways (Level 1, *p* = 0.013) were significantly enriched in C, including metabolic pathways, carbon metabolism, amino acid biosynthesis, fatty acid degradation, and benzoate degradation pathways (level 2). This metabolic enrichment can be linked to the frequent seawater inundation and tidal flushing that C experiences that the ocean supplies diverse organic matter, from labile substrates to recalcitrant compounds such as aromatic hydrocarbons, thereby exerting selective pressure for enhanced heterotrophic degradation capacities (43, 44). In contrast, P exhibited significant enrichment of Cellular Processes (Level 1, *p* = 0.015) and Environmental Information Processing (Level 1, *p* = 0.012), particularly the two-component system and the Pseudomonas aeruginosa biofilm formation pathway (*p* = 0.009). These findings indicate that P communities prioritize signal transduction and biofilm-mediated surface attachment as adaptive strategies, consistent with biofilm colonization patterns in wastewater environments (17, 45–47).

Interestingly, sulfur metabolism did not differ significantly between C and P (*p* = 0.174), despite the presence of SOB and SRB taxa in taxonomic analysis. This taxonomy-function discrepancy likely reflects the low overall abundance of sulfur-cycling bacteria and the inherent limitations of 16S-based functional inference tools like PICRUSt2, which often lack sensitivity for detecting subtle functional shifts (48, 49). Thus, while C communities are metabolically oriented toward organic matter degradation, P communities have evolved a regulatory-and-biofilm-based strategy for persistence, and future metatranscriptomic studies are needed to resolve sulfur-cycling activities.

Co-occurrence network analysis further supported the ecological divergence between C and P. The cross-domain networks (top 15 genera) showed positive correlation ratios of 57.50% in C and 56.18% in P, indicating that both competitive and cooperative interactions coexist. Within-domain networks (top 45 genera) revealed that the bacterial network in C was moderately complex (44 nodes, 199 edges) with a positive co-expression ratio of 68.81%, suggesting balanced competition and cooperation; the fungal network showed an intermediate pattern (positive ratio 89.39%), reflecting the dual potential of fungal mycelia to both compete and cooperate (10, 50); the algal network in P was highly compartmentalised (41 nodes, 74 edges) with a positive ratio of 97.30%, implying predominantly cooperative interactions possibly mediated by shared biofilms (1). Keystone taxa differed between substrates: *Achnanthes*, *Chaetoceros*, *Candida* and *Tunicatimonas* in C versus *Amoeboaphelidium*, *Cloniophora*, *Bisifusarium*, *Tunicatimonas* and *Aspergillus* in P. Keystone species are widely recognised as critical for maintaining network stability and facilitating cross-module communication (2, 48); this difference suggests that distinct ecological processes govern community assembly in the two habitats.

### 4.2 Coexistence of potential corrosion-related but the predominance of net deterioration in service

A major finding of this study is the simultaneous presence of microorganisms potentially involved in microbially induced concrete corrosion (MICC) and concrete healing on both C and P, albeit with different distribution strategies. This coexistence makes the net microbial impact on concrete integrity difficult to predict.

#### 4.2.1 MICC-related taxa exhibit habitat-specific distribution patterns

Among the MICC-associated bacteria (Table 1), acidophilic *Thiobacillus*, which can oxidize H_2_S to H_2_SO_4_ and dissolving cement hydrates to gypsum, occurred in P as a permanently rare but intermediate-distribution taxon. Although its abundance is low, its presence as a “seed bank” may pose a latent corrosion threat if conditions become favorable (2, 3). Sulfate-reducing *Desulfovibrio* and *Desulfobulbus*, which supply H_2_S to SOB, were narrow-distribution rare in C but intermediate-distribution rare in P, reflecting more extensive anaerobic zones in pipes versus localized anaerobic microniches in C (6, 51, 52). The filamentous pioneer SOB *Thiothrix* (narrow in C, intermediate in P) and the fermentative acidogenic *Clostridium* (broad permanently rare in C, intermediate in P) contribute to early neutralization of concrete alkalinity (3, 53, 54). Notably, *Rubrobacter* was the only abundant broad-distribution MICC bacterium that occurred exclusively on C, reflecting its adaptation to salt stress and desiccation on coastal surfaces (38).

MICC-associated Fungi taxa are abundant and widely distributed on C, but differ in P. *Aspergillus*, *Penicillium*, *Cladosporium* and *Trichoderma*, which all secrete strong organic acids (oxalic, citric) that chelate Ca^2+^ and dissolve cement, are abundant and broadly distributed on C, making them key active surface deteriorators (10, 55). In P, the same genera are permanently rare but broadly distributed, likely suppressed by lower oxygen or bacterial competition. *Fusarium*, which penetrates deeply (up to 620 µm) and produces high organic acids, is abundant intermediate in C but abundant broad in P, consistent with its preference for moist, organic-rich pipe environments (14, 55). *Chaetomium* and *Stachybotrys* (narrow in C, intermediate in P) are moisture-dependent cellulolytic fungi associated with damp building materials (11, 35).

MICC-associated Algae taxa are predominantly in coastal concrete. Macroalgae *Rhizoclonium*, *Chaetomorpha* and *Ulva*, which could physically wedge into pores and exude calcium-chelating compounds, are abundant broad or intermediate on C; only *Rhizoclonium* occurs in P (intermediate abundant), as a low-light-tolerant form (12, 13, 56). The unicellular alga *Chlorella* is permanently rare broad in P but intermediate in C, benefiting from stable moisture in pipes (57, 58). Thus, MICC threat in C may be immediate and active from abundant fungi/algae, but in P may be latent and persistent from rare but broadly distributed sulfur-cycling bacteria and diverse rare corrosive fungi.

#### 4.2.2 Many Potential MICP-based healing taxa exhibit in the two habitats

Healing-associated bacteria (Table 2) include well-known MICP agents such as *Bacillus*, like *B. subtilis*, *B. megaterium*, which precipitate CaCO_3_ in cracks. In C, *Bacillus* was permanently rare intermediate-distribution, plus a narrow-distribution rare guild; In P, it was intermediate-distribution rare but more diverse, suggesting a broader “healing seed bank” in pipes (16, 59–61). *Sporosarcina* (including *S. pasteurii*), the model MICP bacterium with extremely high urease activity, was detected in C only as a narrow-distribution rare taxon, possibly limited by lower pH or stronger competition in pipes (18, 59, 62). *Pseudomonas*, which produces EPS-rich biofilms that seal cracks and can also mediate MICP, is more abundant and widespread in P (abundant intermediate) than in C (permanently rare intermediate), reflecting its preference for moist, nutrient-rich pipe environments (43, 63, 64). *Exiguobacterium*, a halophilic MICP bacterium, is abundant and broadly distributed on C but rare intermediate in P, highlighting its adaptation to marine splash zones (9, 65). Actinobacteria include *Arthrobacter*, *Corynebacterium*, *Micrococcus*, and *Rhodococcus* precipitate CaCO_3_ via urease or carbonic anhydrase and produce EPS. In C they are intermediate/narrow-distribution rare, while in P they are more diverse and broadly distributed (66–69). Interestingly, halotolerant genera (*Halomonas*, *Halobacillus*, *Bhargavaea*, *Chryseomicrobium*) are as intermediate-distribution rare in P, but as narrow-distribution rare in C, indicating they may be more competitive in the stable, slightly saline pipe environment (67, 69–71). *Paenibacillus* is narrow in C but broad in P, indicating broader ecological tolerance in pipes (72, 73).

Fungal healing taxa showed distinct patterns. *Aspergillus* (abundant broad in both C and P) precipitates CaCO_3_ via urease or carbonic anhydrase, with hyphae acting as nucleation sites. *Penicillium* and *Trichoderma*, both capable of MICP and EPS secretion, are abundant broad on C but only permanently rare broad in P, suggesting that they are stronger competitive in dynamic coastal environment (21, 74–76). The yeasts *Candida* and *Rhodotorula*, and *Fusarium*, which use non-ureolytic pathways, e.g., acetate-based, to precipitate CaCO_3_ without ammonia release, are abundant intermediate -distribution on C but most abundant broad -distribution in P, indicating that pipes provide a more favourable nutrient base for them (20, 23, 77). *Neurospora*, urease-positive, forms biomineralized mycelial networks, and the yeast *Naganishia*, precipitates vaterite, are detected exclusively in P as permanently rare intermediate taxa, likely due to specific organic nutrient requirements met only in pipe environments (23, 78, 79).

#### 4.2.3 Functional duality within genera but the predominance of net deterioration in service

An important complexity is that both potential MICC-related and MICP-based healing taxa exist simultaneously in C and P (Table 1-2), and many genera even contain both corrosion-promoting and healing-promoting species. For example, *Aspergillus* includes corrosive species (*A. niger*, *A. fumigatus*, *A. versicolor*, etc.) that secrete organic acids and degrade concrete (35, 80, 81), as well as healing species (*A. nidulans*, *A. iizukae*, and *A. flavus*) that precipitate CaCO_3_ healing the cracks of concrete (22, 82, 83). Similar duality exists in *Penicillium*, *Fusarium*, *Rhodotorula*, *Bacillus*, *Pseudomonas*, and many other genera. However, despite the co-occurrence of potentially healing taxa, concrete infrastructures invariably undergo net deterioration over their service life under natural conditions.

This predominance of corrosion over healing arises from several factors. First, the environmental conditions of most concrete surfaces are not conducive to the enrichment of healing-related microorganisms. In our study, although both C and P harbored MICP-related taxa (Table 2), most of these healing bacteria were classified as permanently rare (relative abundance always ≤1%), especially in P, where they constitute a latent “seed bank” rather than an active, abundant community. Second, the physiological requirements for effective MICP such as sufficient calcium ions, suitable pH, and carbon source, are not always met in unamended concrete, whereas the metabolic pathways driving corrosion, such as organic acid production and sulfur oxidation, are directly fueled by the ambient environment. Third, even when healing taxa are present, their remedial capacity is typically localized and slow, whereas corrosion processes can propagate rapidly through chemical and physical feedback loops, such as acidification exposing fresh surface.

## 5 Conclusions

This study provides a comprehensive comparative analysis of bacterial, fungal, and algal communities on coastal concrete (C) and concrete drainage pipes (P). Fungal and algal α-diversity are significantly higher in P than in C, whereas bacterial diversity does not differ significantly; β-diversity reveals strong separation of bacterial and algal communities between the two habitats, but not for fungi. A shared core “seed bank” of 575 bacterial, 520 fungal, and 40 algal ASVs exists across both substrates, suggesting a ubiquitous microbial reservoir in coastal concrete environments. Environmental filtering by salinity, NH_4_^+^-N, NO_3_^-^-N, and COD drives community differentiation that pipes with low salinity, high nutrients enrich oligotrophic degraders *Sphingomonas* and saprotrophic/acid-producing fungi like *Arxiella* and *Bisifusarium*, whereas coastal concrete (high salinity, low nutrients) selects for halotolerant EPS-producing bacteria (Tunicatimonas, Muricauda) and the extremotolerant alga Coelastrella. Functional prediction indicates that bacterial communities in C favor metabolic pathways, while those in P prioritize environmental signal transduction and biofilm formation. Co-occurrence networks show distinct keystone taxa and higher positive correlations within fungi and algae, indicating domain-specific ecological strategies. Notably, microorganisms potentially involved in microbially induced concrete corrosion and those capable of concrete healing coexist on both substrates but exhibit different distribution patterns. Despite this co-occurrence of healing potential, concrete deterioration remains the net outcome under natural conditions due to the low abundance of active healers, unfavorable environmental conditions for microbial healing, and rapid feedback loops that accelerate corrosion. This study establishes a functional taxonomic framework for understanding the dual roles of concrete-associated microbiomes and provides a basis for developing microbiome-based durability management strategies, such as activating the rare healing reservoir in pipes or promoting endogenous healing taxa in coastal structures.

## Disclosure statement

The authors declare no conflict of interest.

## Funding

This research was funded by the Guangxi Science and Technology Program (Guike ZY24212001 and Guike XT2503960022)

This research was funded by the Guangxi Science and Technology Program (Guike XT2503960022).

